## Supplementary material for "A surface morphology-based inference method for the cell wall elasticity profile in tip-growing cells": S1 Text

### Supplemental Figures

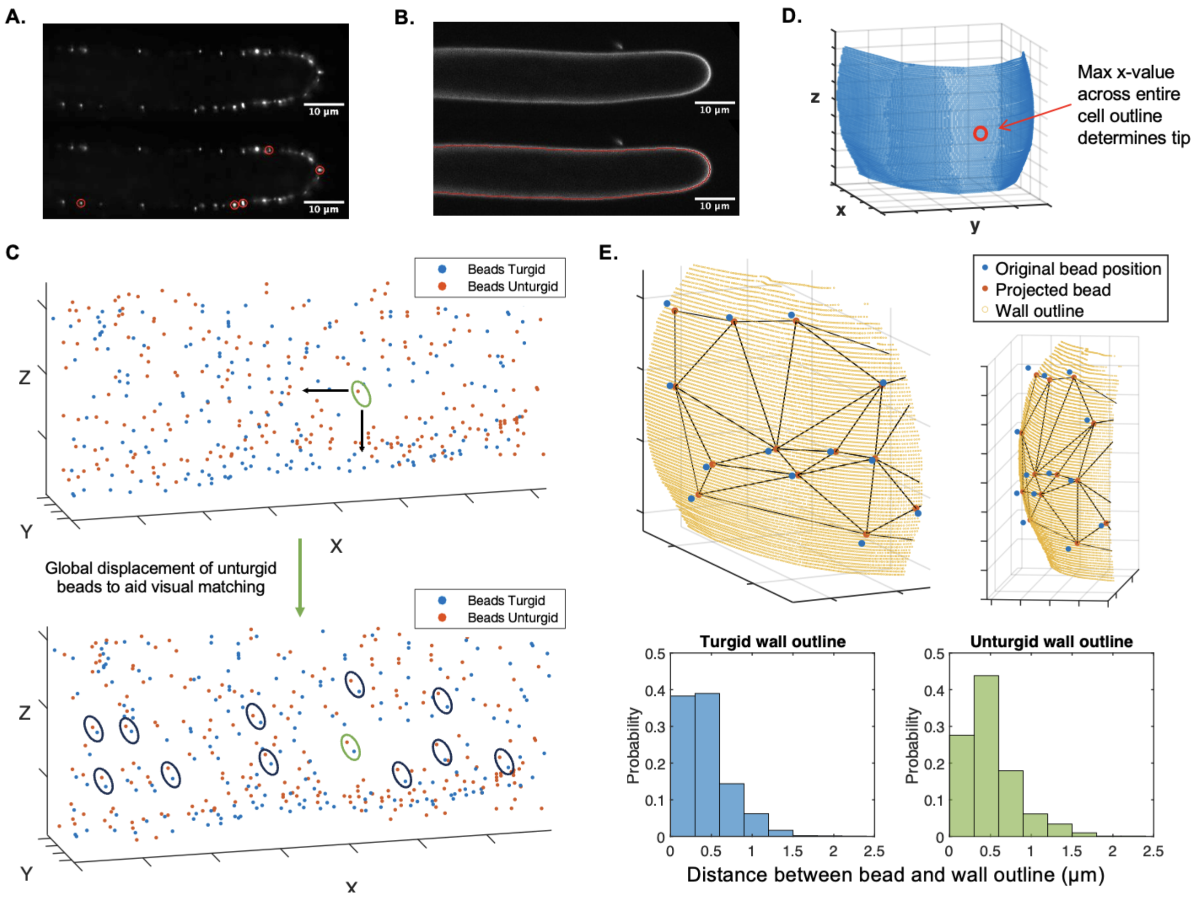


**Fig S1: Conversion of experimental data into MATLAB and inference method details. A.** Example of marker point localization in ImageJ. Sub-pixel resolution locations of fluorescent points are identified by the RS-FISH plugin. **B.** Example of cell wall outline localization using the Ridge Detection plugin in ImageJ. **C.** Demonstration of marker point matching between the two configurations in MATLAB. Matching is done manually through visual confirmation of the two sets of marker points. Additional global displacement of one set of marker points may be needed to aid in the visual matching (bottom). **D.** Identification of tip point is done using the wall outline data and the max x-value. **E.** Demonstration of marker point projection from two views (top). Histogram depicting distances between the bead and wall outline for both the turgid and unturgid wall outlines (bottom).

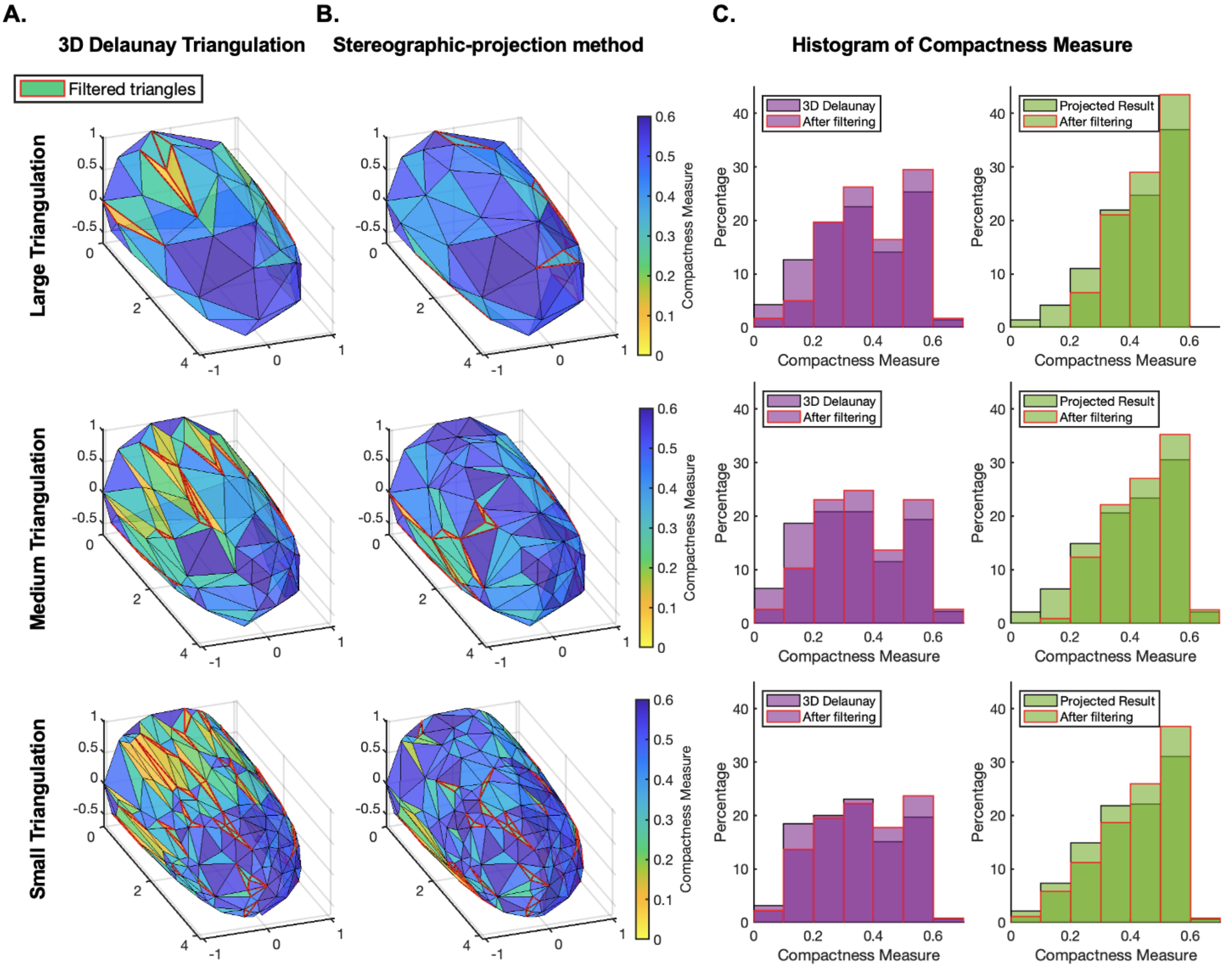


**Fig S2: Comparison of triangulation techniques. A.** Surface triangulation using 3D Delaunay algorithm. **B.** Stereographic-projection method presented in our main text, on the same set of marker points. For both **A.** and **B.**, the triangle color indicates the value of its compactness score: $4\pi\text{Area}/\text{Perimeter}^2$. The triangles outlined in red are triangles filtered out using our sensitivity analysis algorithm. **C.** Histogram of compactness score for the two triangulations. The histogram outline in red is the result after filtering out the triangles shown in red in **A.** and **B.** respectively.

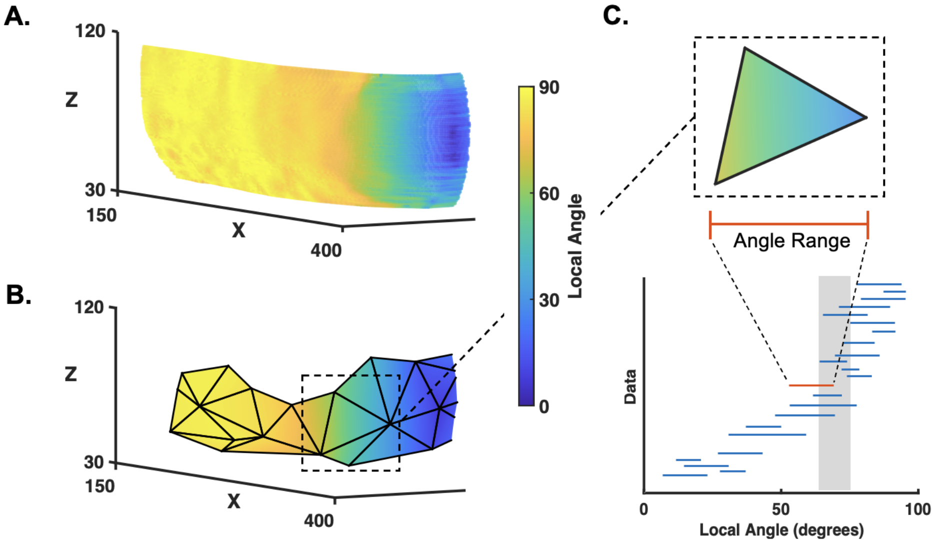


**Fig S3: Defining local angle on the wall surface points and triangulation.** **A.** Local angle of each cell wall outline point calculated using $\alpha = \cos^{-1}(\hat{n} \cdot \hat{z})$. **B.** Triangulation map of local angle. **C.** (Top) Example triangle with local angle range. (Bottom) Example mapping of data along the local angle. Each triangle is represented by a line showing its angle range, and then it is sorted into its bins (example bin in gray).

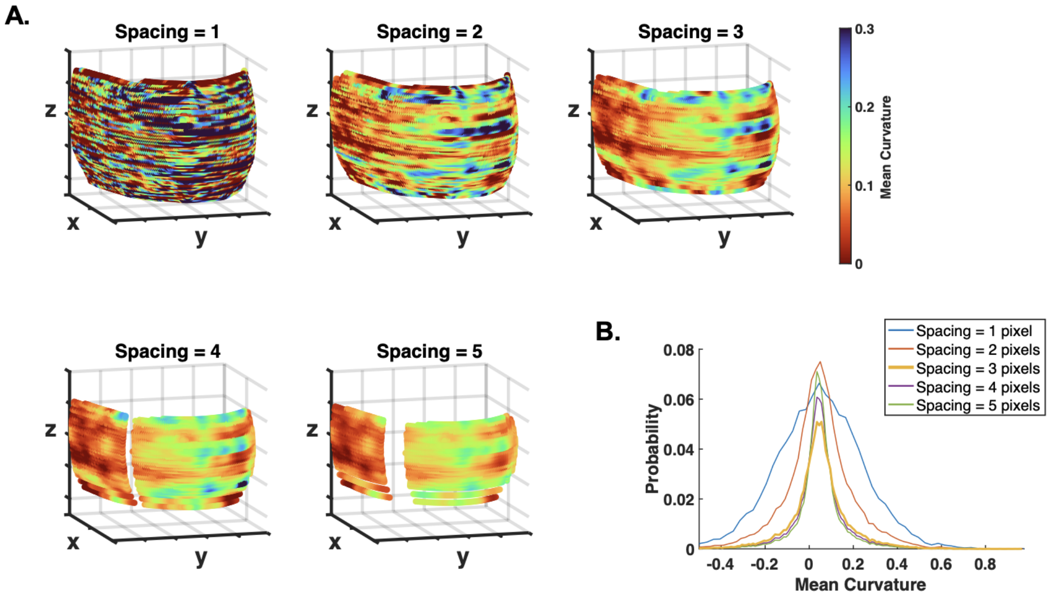


**Fig S4: Confirmation of chosen spacing size in curvature calculation.** **A.** Mean curvature results at 5 different spacing levels shown. **B.** The corresponding probabilities of mean curvature at the 5 spacing levels. We have chosen the bolded yellow line, spacing = 3 pixels.

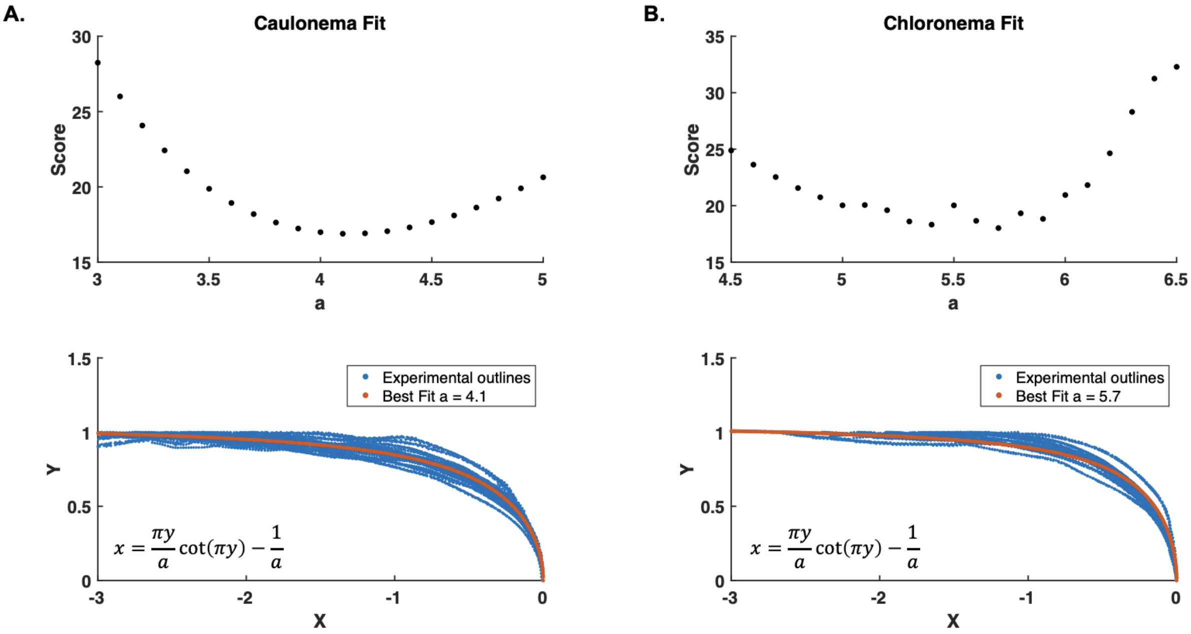


**Fig S5: Experimental caulonema and chloronema hyphoid parameter fitting. A.** Caulonema fitting, best fit at $a = 4.1$. **B.** Chloronema fitting, best fit at $a = 5.7$. (Top) Score of fit versus the parameter $a$ shown in the equation below. (Bottom) Best fit hyphoid curve shown in red on top of 6 experimental cell outlines (both sides of each cell outline are included).

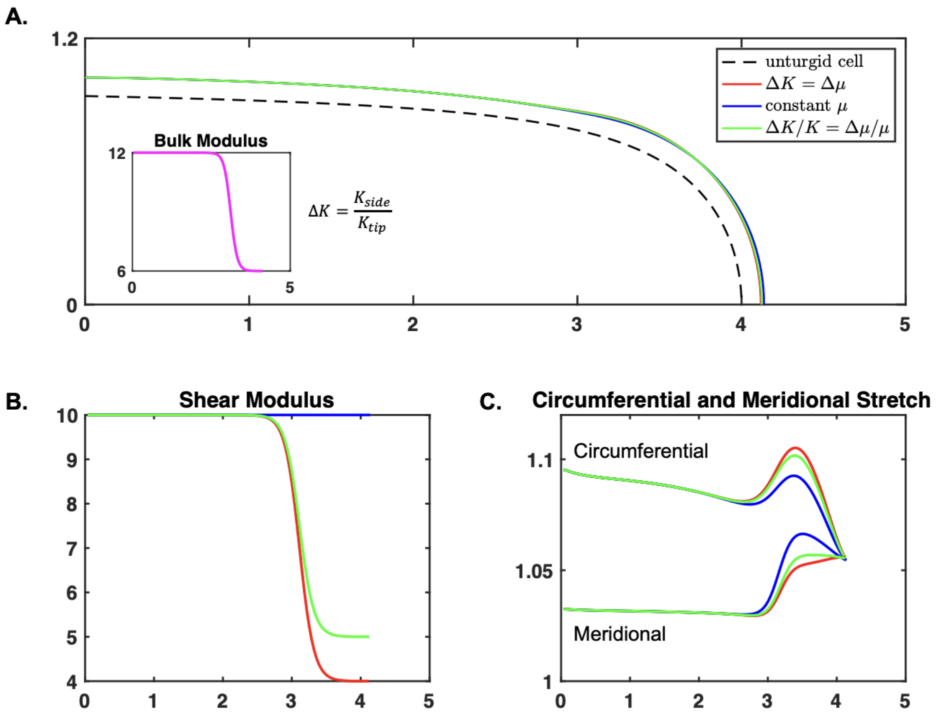


**Fig S6: Effect of different $\mu_h$ gradients in relation to the $K_h$ gradient** Three gradient cases are studied: the ratio of the side to the tip are equal for both $K_h$ and $\mu_h$, $\mu_h$ is constant, and the relative ratio of $K_h$ and $\mu_h$ is equal. **A.** The turgid cell shapes of the three cases are compared. Inset shows the $K_h$ gradient. **B.** The respective $\mu_h$ gradients shown. **C.** The respective elastic stretch ratios for the three cases shown.

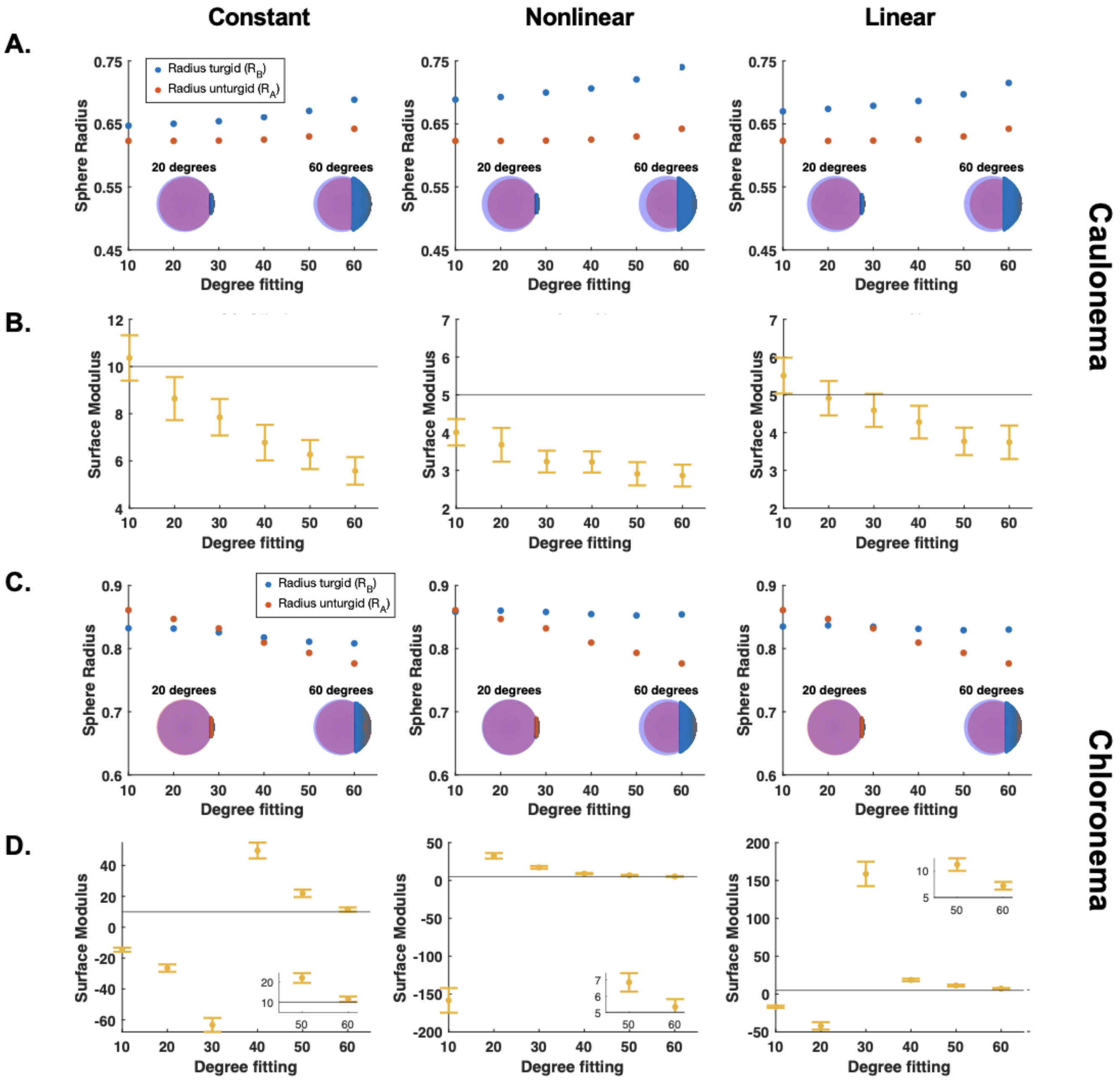


**Fig S7: Sphere-fitting method to measure synthetic caulonema and chloronema tip surface modulus.** **A.** Sphere radius of the turgid (blue) and unturgid sphere fitting (red) versus the maximum degrees of the synthetic cell outline used to fit the sphere. The three gradient cases are shown for each cell type. Visual sphere fit result examples are shown in the inset. **B.** Corresponding modulus value calculated by $Y_{tip} = \frac{R_B}{2(R_B - R_A)/R_A}$ where $R_A$ is the unturgid sphere radius and $R_B$ is the turgid sphere radius. Insets on some graphs show re-focused view. Chloronema results are shown in **C.** and **D.**. Modulus values are non-dimensionalized.

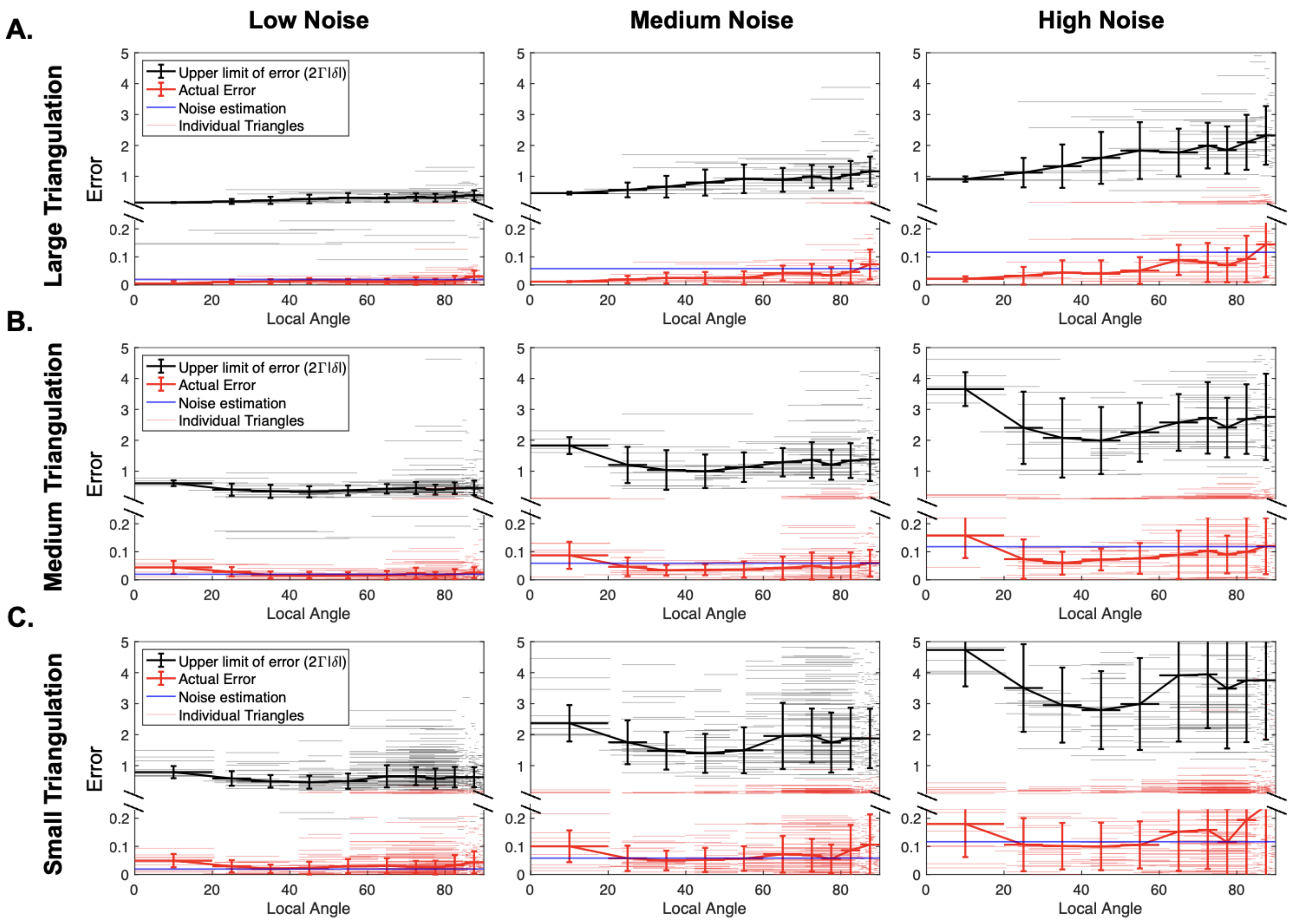


**Fig S8: Area stretch ratio error is below theoretical limit and comparable to displacement noise.** For the three triangulation cases **A.** large, **B.** medium, and **C.** small, the relative error of $\lambda_1\lambda_2$ is shown in red. The theoretical upper limit of the error using the function $2\Gamma|\delta|$ is shown in black. For both values, the individual triangles and binned average are plotted (see legend). For each noise case, we estimated the upper limit of the displacement noise $|\delta|$ (blue line).

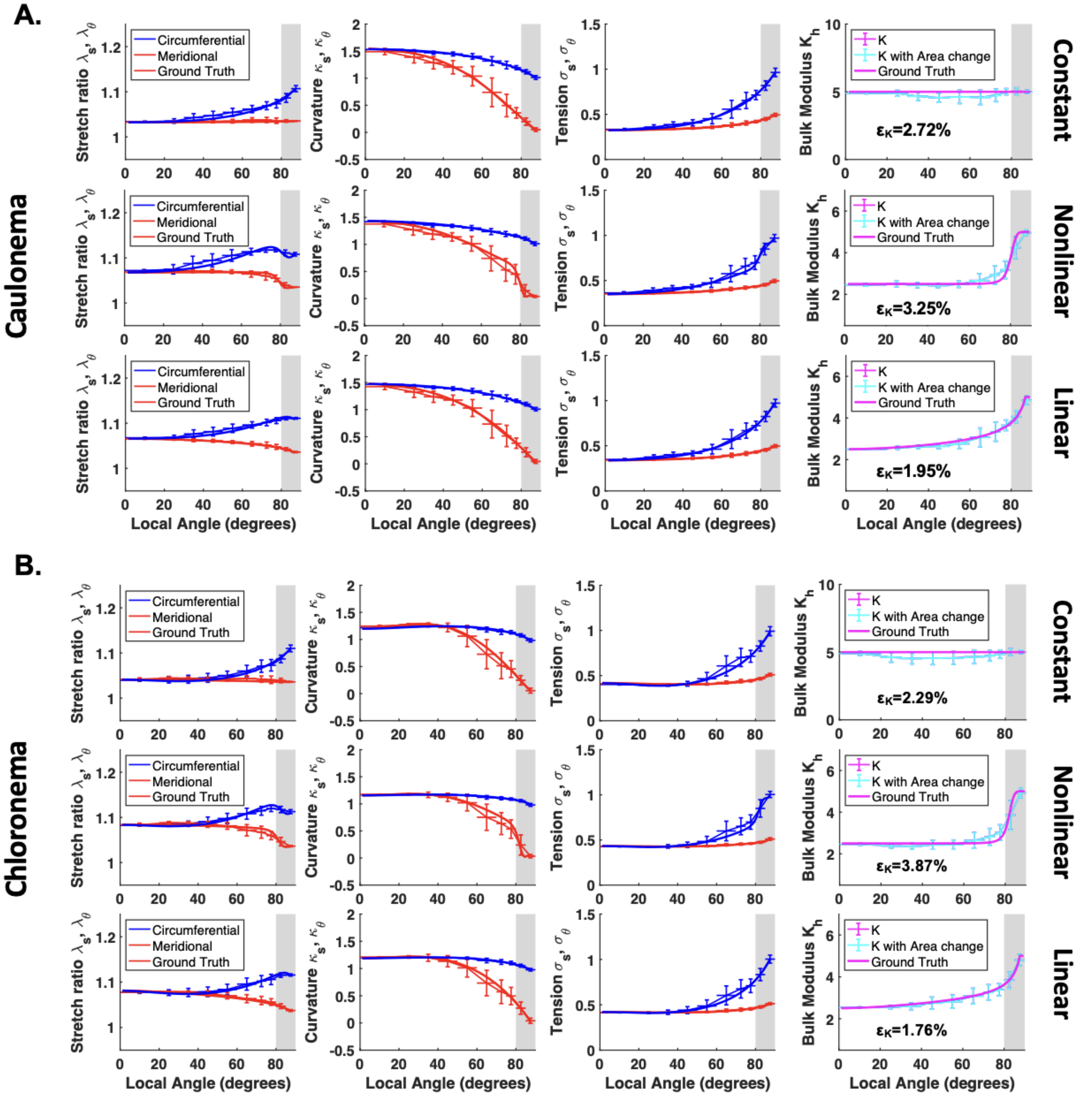


**Fig S9: No noise inference of geometric parameters in synthetic caulonema and chloronema cells.** Inference results of a single **A.** caulonema and **B.** chloronema cell with no noise. Each part shows an inference on cell's with a constant, nonlinear, and linear distributions. In each case, the elastic stretch ratio, curvature, and tension inferences are shown with their corresponding ground truth values. The rightmost panels depict the bulk modulus inference along with their relative error from the ground truth, $\varepsilon_K$. The cyan curve depicts the inference of $K_h$ calculated with $S_B/S_A$. Note that in the linear case, the curve will be linear when plotted as a function of $z$, instead of $\alpha$. Values are non-dimensionalized ($\kappa_{s,\theta} = \bar{\kappa}_{s,\theta}\bar{L}$, $\sigma_{s,\theta} = \bar{\sigma}_{s,\theta}/\bar{P}\bar{L}$, $K_h = \bar{K}_h/\bar{P}\bar{L}$).

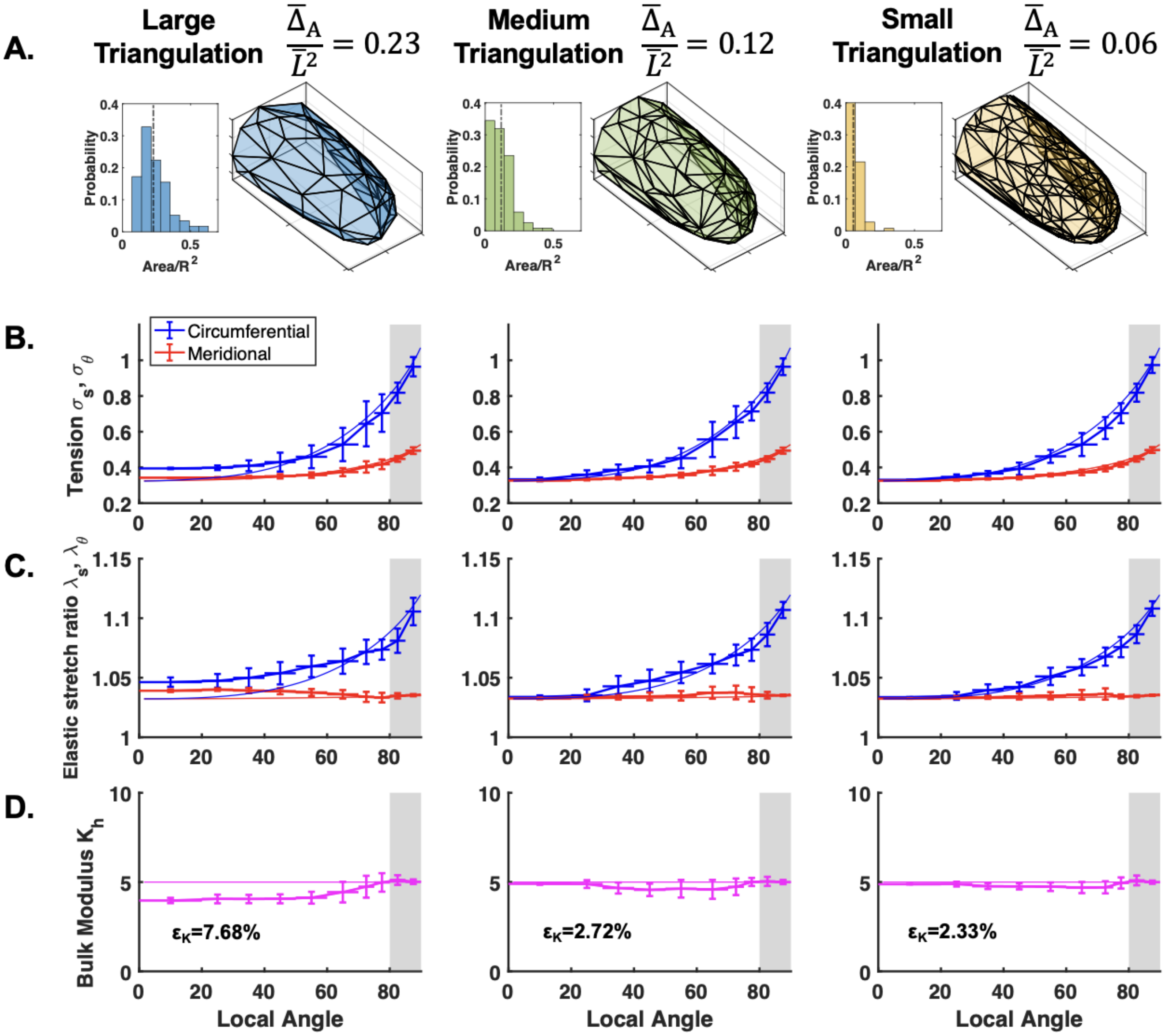


**Fig S10: Triangulation size analysis with no noise. A.** Representative triangulation on a synthetic caulonema cell and histogram of normalized triangle areas. Dashed line on histogram represents mean area displayed above to define each triangulation size: $\bar{\Delta}_A/\bar{L}^2$. **B.** Corresponding tension results from the three different triangulations. **C.** Corresponding elastic stretch ratio results from the three different triangulations. **D.** Corresponding bulk modulus results from the three different triangulations. The average relative error ($\varepsilon_K$) of the inference is shown. Values are non-dimensionalized ($\sigma_{s,\theta} = \bar{\sigma}_{s,\theta}/\bar{P}\bar{L}$, $K_h = \bar{K}_h/\bar{P}\bar{L}$).

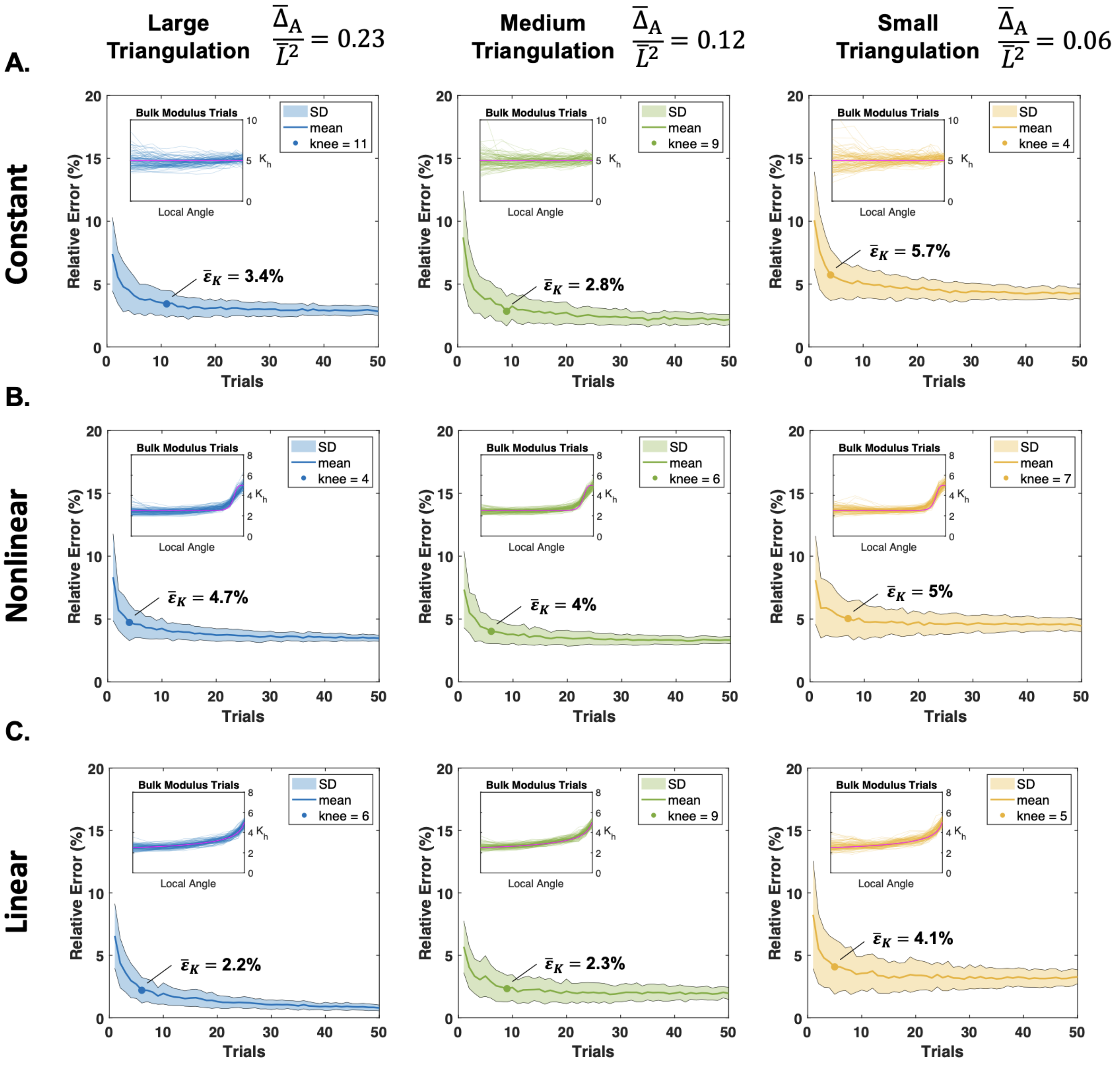


**Fig S11: Low noise caulonema trial analysis.** Average relative error of the bulk modulus inference for **A.** constant, **B.** nonlinear, and **C.** linear gradients versus the number of trials for three triangulation cases. In each case, $n$ trials are chosen from 100 total cells, and this is repeated 100 times to obtain the standard deviation and mean shown in each graph. The corresponding 100 cell rescaled bulk modulus inferences are shown in each inset on top of the ground truth bulk modulus in magenta. Additionally, the knee of the curve is plotted along the mean, with its averaged error ($\bar{\varepsilon}_K$) at that trial number.

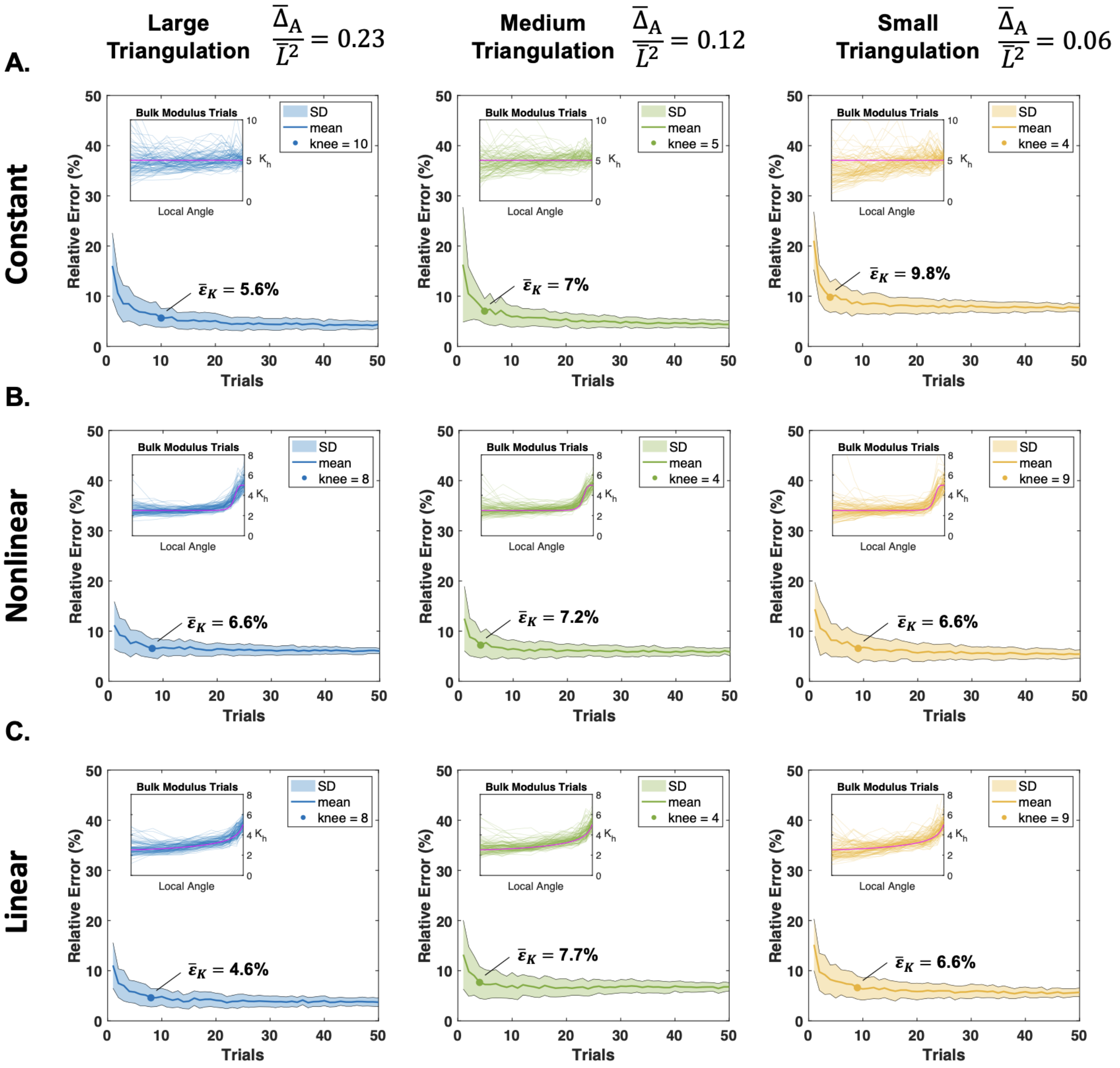


**Fig S12: Medium noise caulonema trial analysis.** Average relative error of the bulk modulus inference for **A.** constant, **B.** nonlinear, and **C.** linear gradients versus the number of trials for three triangulation cases. In each case, $n$ trials are chosen from 100 total cells, and this is repeated 100 times to obtain the standard deviation and mean shown in each graph. The corresponding 100 cell rescaled bulk modulus inferences are shown in each inset on top of the ground truth bulk modulus in magenta. Additionally, the knee of the curve is plotted along the mean, with its averaged error $(\bar{\varepsilon}_K)$ at that trial number.

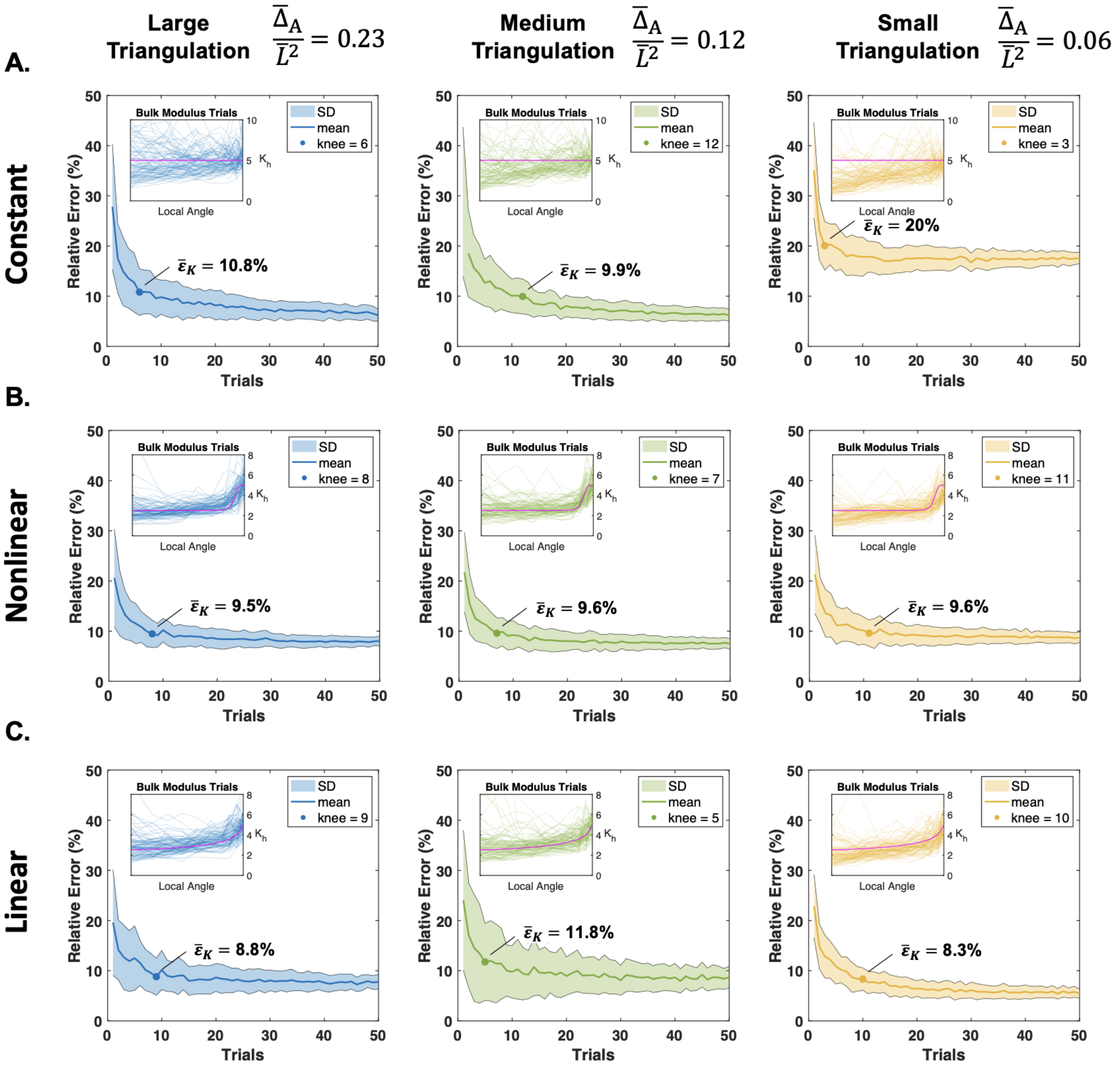


**Fig S13: High noise caulonema trial analysis.** Average relative error of the bulk modulus inference for **A.** constant, **B.** nonlinear, and **C.** linear gradients versus the number of trials for three triangulation cases. In each case, $n$ trials are chosen from 100 total cells, and this is repeated 100 times to obtain the standard deviation and mean shown in each graph. The corresponding 100 cell rescaled bulk modulus inferences are shown in each inset on top of the ground truth bulk modulus in magenta. Additionally, the knee of the curve is plotted along the mean, with its averaged error ($\bar{\varepsilon}_K$) at that trial number.

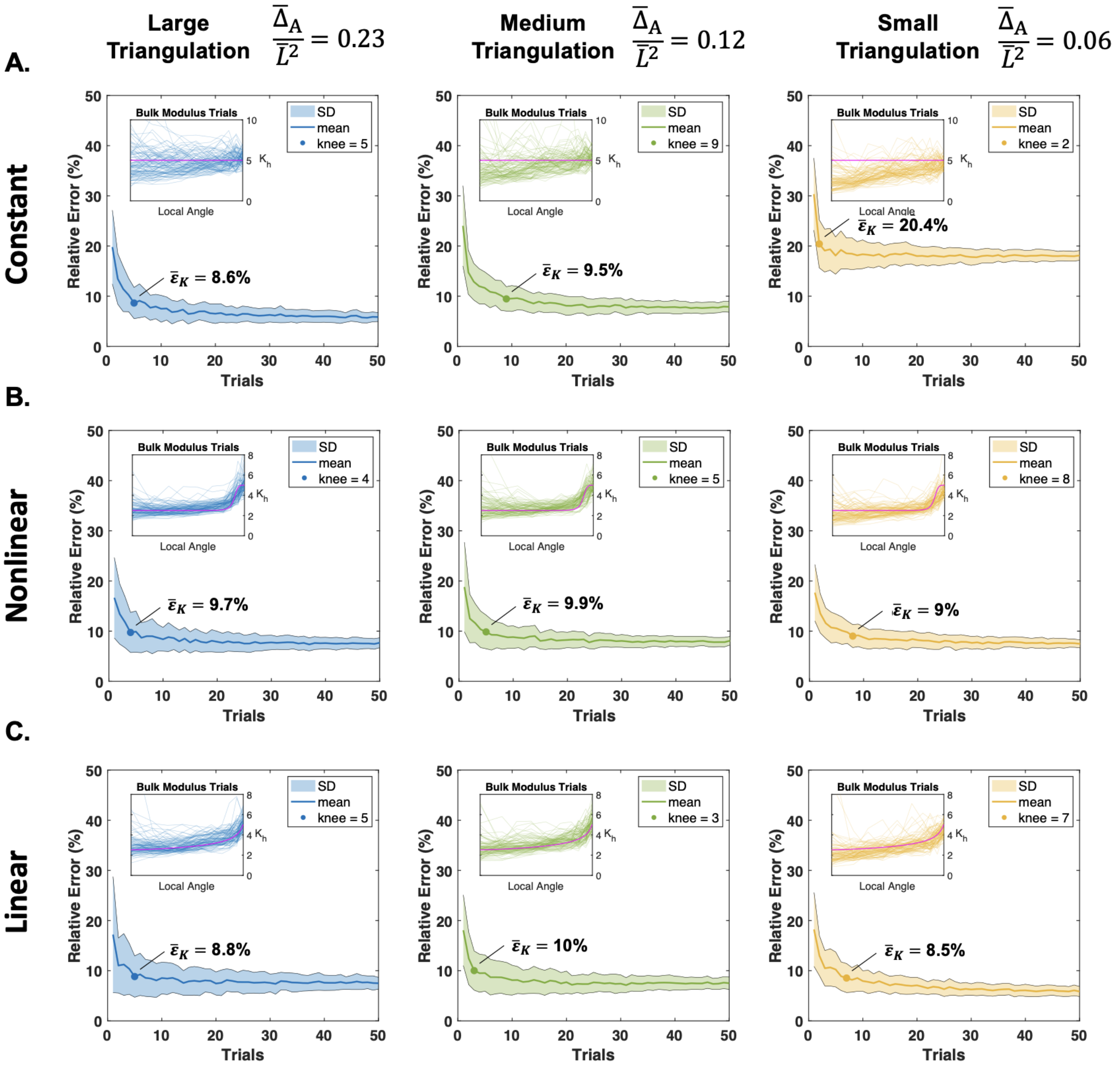


**Fig S14: High noise (two triangulation) caulonema trial analysis.** Average relative error of the bulk modulus inference for **A.** constant, **B.** nonlinear, and **C.** linear gradients versus the number of trials for three triangulation cases. In each case, $n$ trials are chosen from 100 total cells (with two triangulations each, see S1 Text), and this is repeated 100 times to obtain the standard deviation and mean shown in each graph. The corresponding 100 cell rescaled bulk modulus inferences are shown in each inset on top of the ground truth bulk modulus in magenta. Additionally, the knee of the curve is plotted along the mean, with its averaged error ($\bar{\varepsilon}_K$) at that trial number.

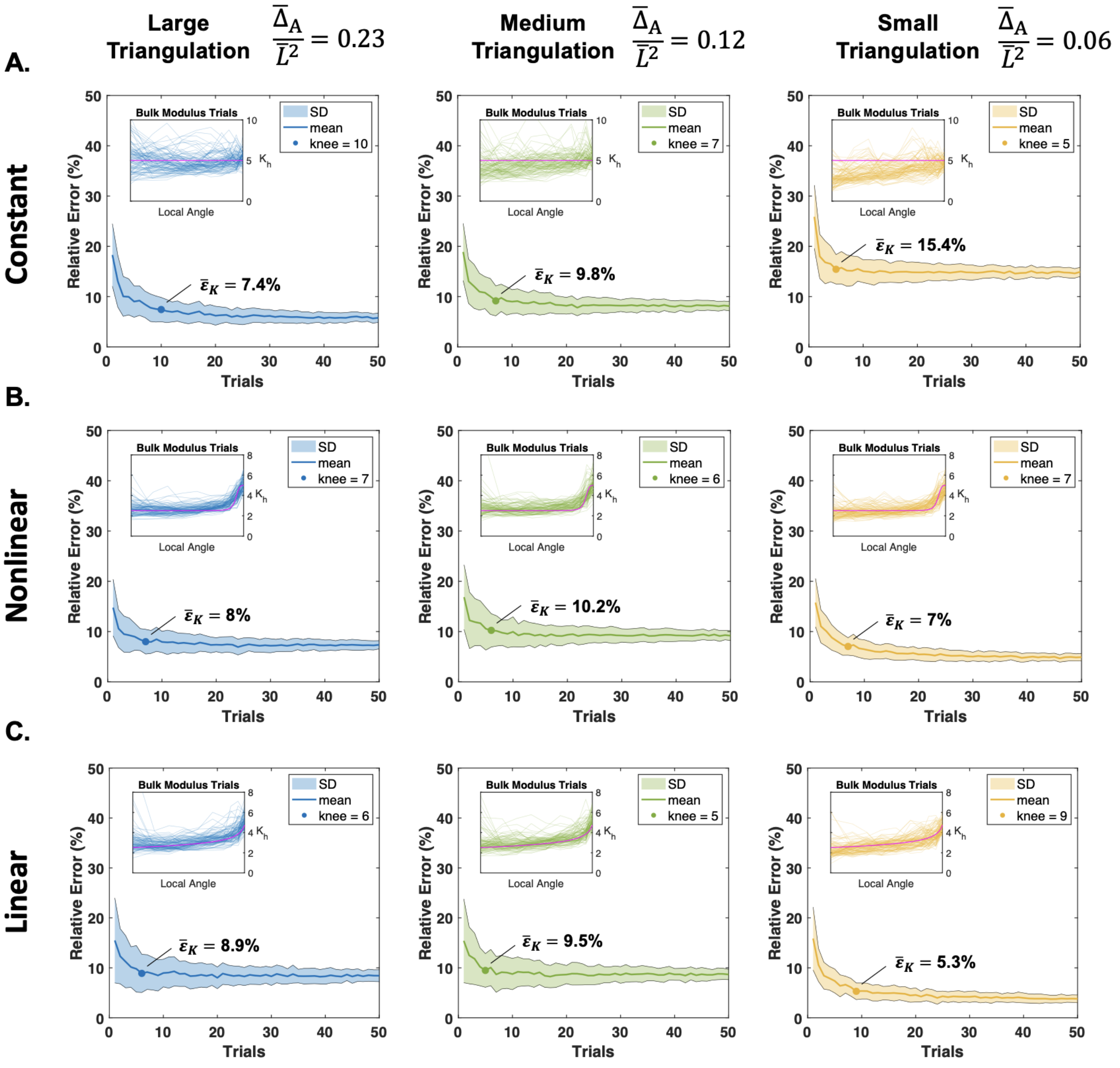


**Fig S15: High noise (two triangulation) chloronema trial analysis.** Average relative error of the bulk modulus inference for **A.** constant, **B.** nonlinear, and **C.** linear gradients versus the number of trials for three triangulation cases. In each case, $n$ trials are chosen from 100 total cells (with two triangulations each, see S1 Text), and this is repeated 100 times to obtain the standard deviation and mean shown in each graph. The corresponding 100 cell rescaled bulk modulus inferences are shown in each inset on top of the ground truth bulk modulus in magenta. Additionally, the knee of the curve is plotted along the mean, with its averaged error ($\bar{\varepsilon}_K$) at that trial number.

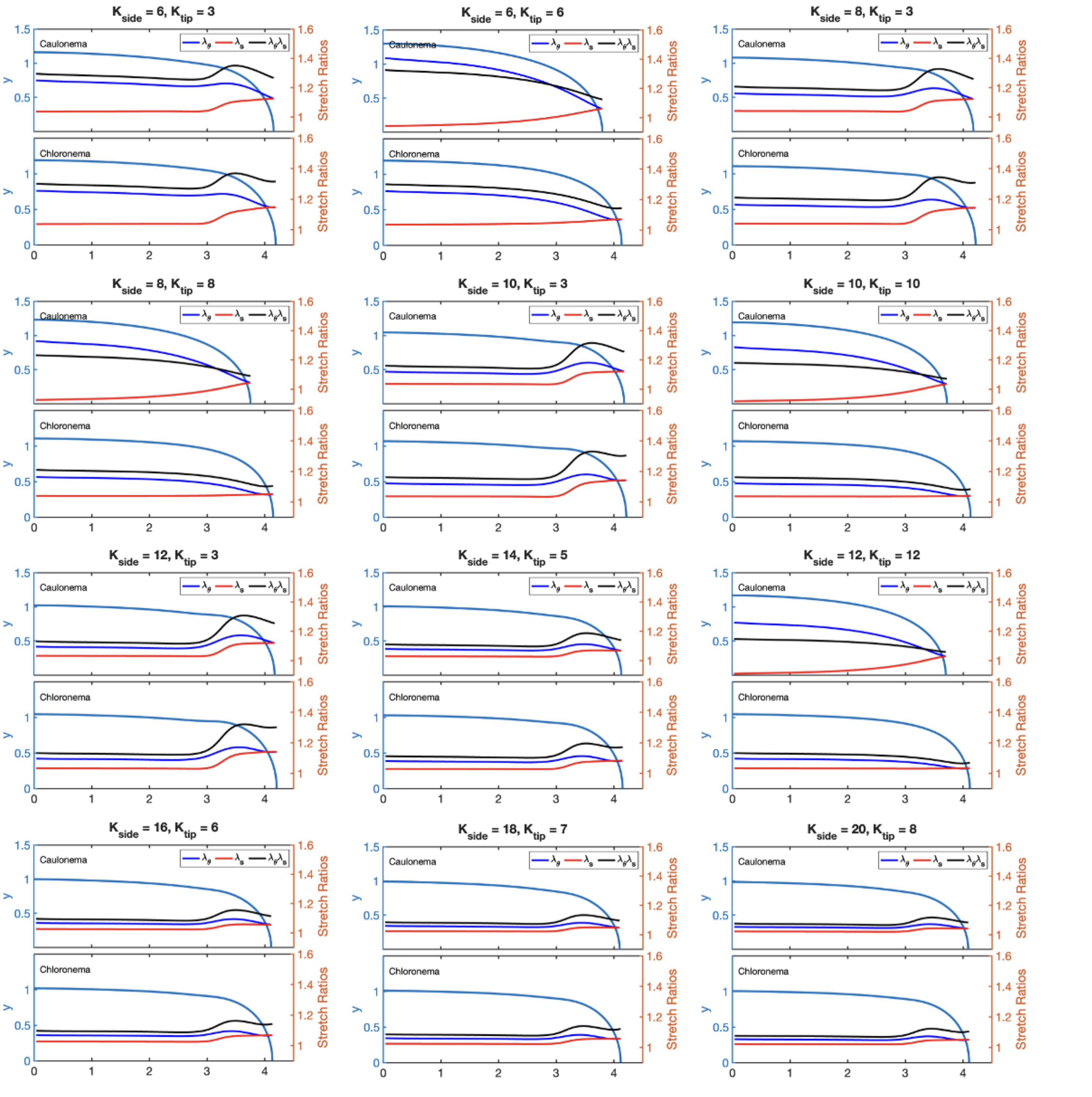


**Fig S16: Subset of caulonema and chloronema cell outlines and elastic stretch ratio.** For each subplot, the turgid cell outlines of the respective caulonema and chloronema cells are plotted (left axis). The circumferential (blue), meridional (red), and area stretch ratio (black) are plotted on top (right axis).

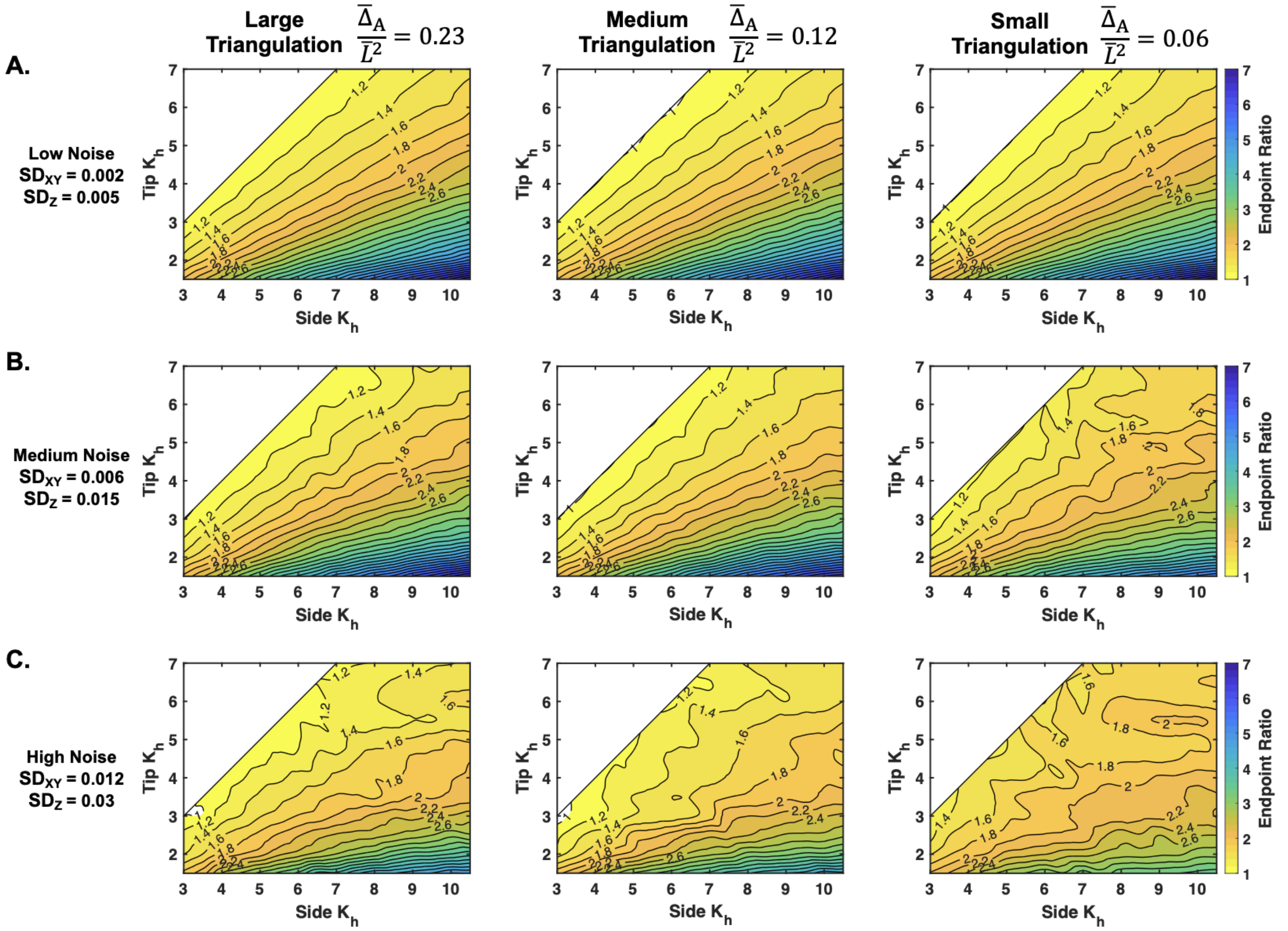


**Fig S17: Inferred endpoint ratio results across different triangulation sizes and noise levels from a synthetic chloronema cell.** Contour map depicts the inferred endpoint ratio levels ($K_{side}/K_{tip}$) across a range of $K_{side}$ and $K_{tip}$ values at the three triangulation sizes and different noise levels: **A.** low noise, **B.** medium noise, and **C.** high noise. Results come from repeating the averaging of the standardized 10 cells to remove random effects (see S1 Text). Modulus values are non-dimensionalized ($K_h = \bar{K}_h/\bar{P}\bar{L}$).

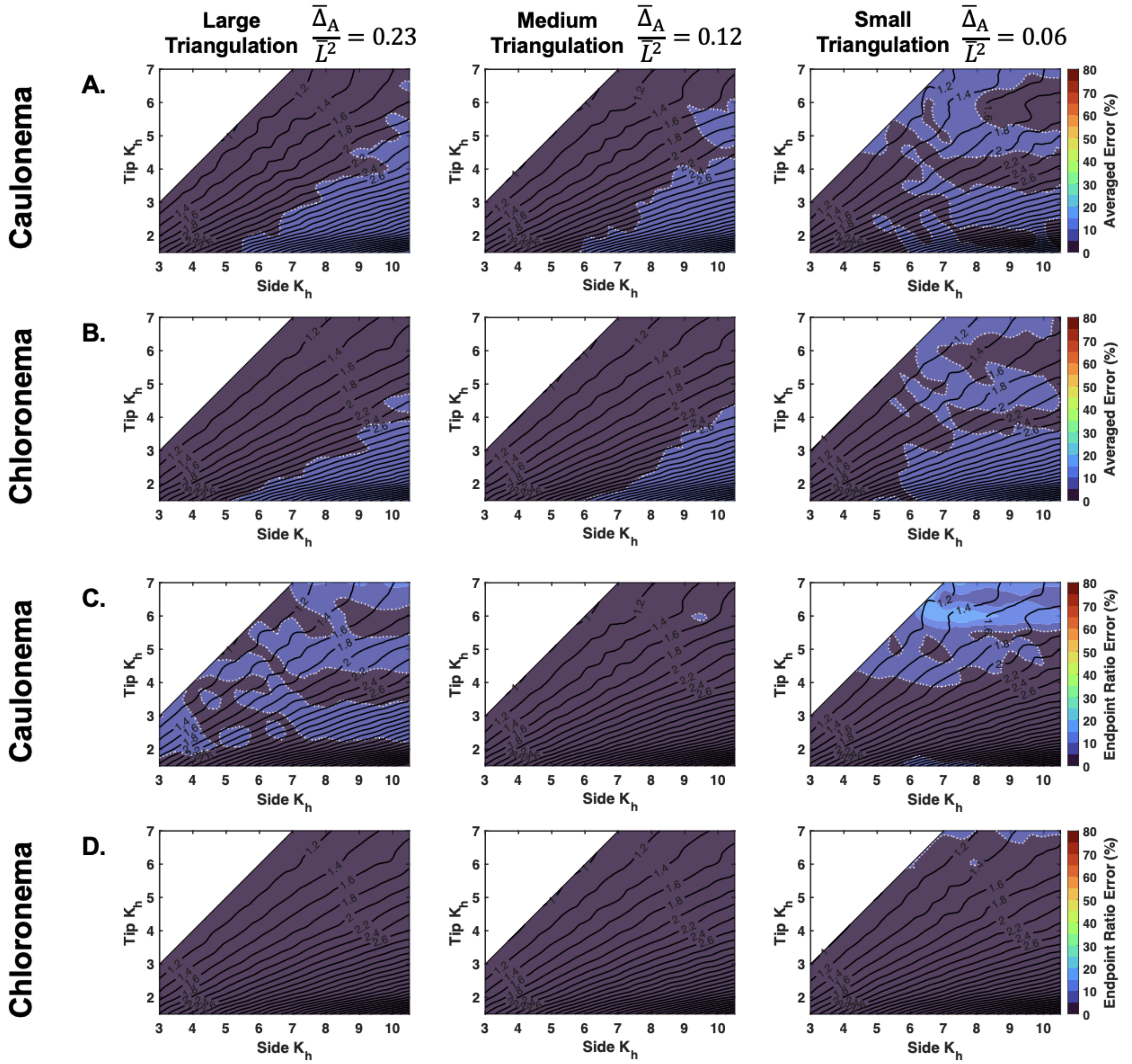


**Fig S18: Two error quantifications for caulonema and chloronema cell bulk modulus inference with low noise** Averaged error ($\bar{\varepsilon}_K$) for **A.** caulonema and **B.** chloronema cells. Relative absolute value endpoint ratio error ($\bar{\varepsilon}_{K_{side}/K_{tip}}$) for **C.** caulonema and **D.** chloronema cells. In all parts, the underlying contours from Fig 7 (caulonema) and Fig S17 (chloronema) are shown. The white dashed line denotes the boundary of 20% error and the white dotted line denotes the boundary of 5% error. Results come from repeating the averaging of the standardized 10 cells to remove random effects (see S1 Text). Modulus values are non-dimensionalized ($K_h = \bar{K}_h/\bar{P}\bar{L}$).

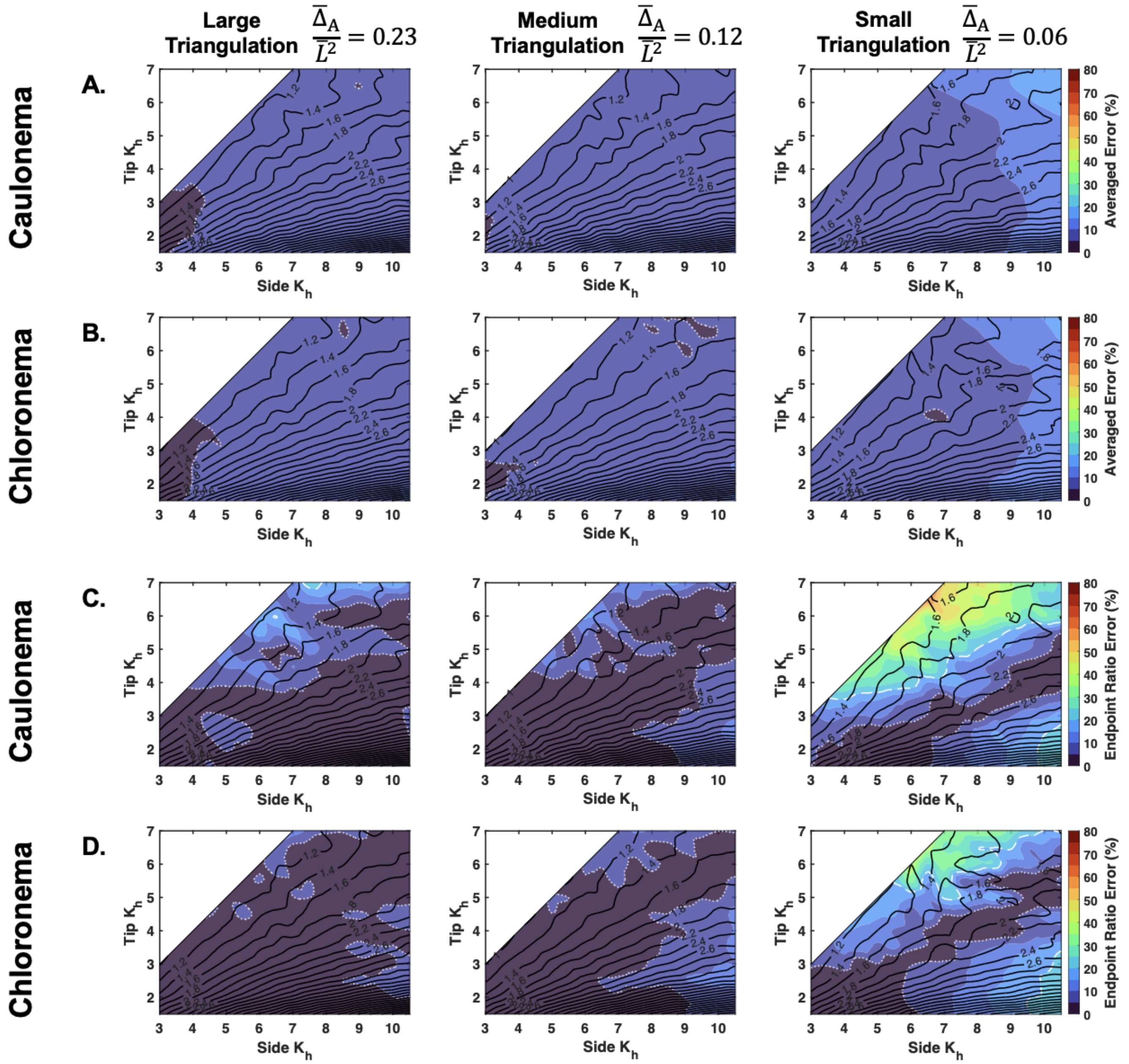


**Fig S19: Two error quantifications for caulonema and chloronema cell bulk modulus inference with medium noise** Averaged error ($\bar{\varepsilon}_K$) for **A.** caulonema and **B.** chloronema cells. Relative absolute value endpoint ratio error ($\bar{\varepsilon}_{K_{side}/K_{tip}}$) for **C.** caulonema and **D.** chloronema cells. In all parts, the underlying contours from Fig 7 (caulonema) and Fig S17 (chloronema) are shown. The white dashed line denotes the boundary of 20% error and the white dotted line denotes the boundary of 5% error. Results come from repeating the averaging of the standardized 10 cells to remove random effects (see S1 Text). Modulus values are non-dimensionalized ($K_h = \bar{K}_h/\bar{P}\bar{L}$).
