## Supplemental Figures for "A surface morphology-based inference method for the cell wall elasticity profile in tip-growing cells"

### Contents

1.1 Experimental protocols and moss imaging
1.2 Wall surface data processing, and marker point matching and projection:
1.3 Computation of curvatures from the wall surface:
1.4 Triangle error analysis
1.5 Experiment noise estimation
1.6 Range and distribution of elastic moduli inputs to generate synthetic cells:
1.7 Data integration and multiple cell averaging

#### 1.1 Experimental protocols and moss imaging

All preliminary experiments used the wildtype line (Gransden) of *P. patens* cultured over cellophane disks in WPi solid media (Macronutrients: 1 mM MgSO4, 1mM Ca(NO3)2, 4 mM KNO3, 89 $\mu$M Fe-EDTA, 1.84 mM KH2PO4. Micronutrient details can be found in [1]. Media was adjusted to pH 5.5. Plants were grown at 25°C with a cycle of 8 hours dark and 16 hours light. To obtain the protonema cells, we ground the plants and sieved them through a 70 $\mu$m Nylon cell strainer (BD Falcon) before replating them. After 7 days, the plants were resubmerged into liquid WPi media before being transferred to our microfluidic devices for imaging.

Two types of 0.2 µm fluorescent beads were used in the experiments: Red FluoSpheres amine-modified and yellow-green FluoSpheres sulfate-modified (ThermoFisher). Both were prepared in the same way. The bead preparation protocol was adapted from [2]. Each time, a 1:2000 dilution of beads (2% solid stock) in moss media was made. This was then bath sonicated for 1 minute and centrifuged at 15000 rpm for 3 minutes. Calcofluor white dye was used when the yellow-green FluoSpheres were used, and Direct Yellow 96 dye was used with the red FluoSpheres. In both cases, a final concentration of 1 µg/mL was used and 1 mL of the fluorescent bead solution was added through liquid flow in a microfluidic device.

Images seen in Fig 1 and 2 were taken with a Nikon Ti-E microscope equipped with a Yokogawa CSU-X1 spinning disk confocal head using a 40X oil-immersion objective (S Fluor N.A. 1.30). The yellow-green beads and Direct Yellow dye were imaged with the 488 nm laser. The red beads were imaged with the 561 nm laser, and the calcofluor white dye was imaged with a 402 nm laser. Images were acquired with the Molecular Devices MetaMorph v7.7 software and Hamamatsu Flash 4.0 LT sCMOS camera. A Z-Stack step-size of 0.16 µm was used to match the dimensions of the XY directions, with a total Z-range of 15-20 µm to cover the entire cell.

#### 1.2 Wall surface data processing, and marker point matching and projection:

To see the settings we used to extract the raw data, as well as a full demonstration of the processes described in this section, please refer to our GitHub repository: `https://github.com/rholee-xu/surface-model/tree/main`.

**Marker point and wall outline localization:** Images were opened and processed in ImageJ. To pinpoint the location of the fluorescent beads, the plug-in RS-FISH was used [3]. To extract the wall outline, the plugin Ridge Detection was used [4, 5]. Besides the line width, in most cases the default settings were used. We used a line width of 9-12 for our cell wall outline. When there was noise in the image, the contrast, sigma, and contrast parameters were adjusted to obtain the best result visually. The same settings were run on every slice of the stack.

**Marker point subsetting and matching:** To select and match the marker point locations between the two configurations, we superimposed the bead positions in the turgid and unturgid configurations (Fig S1C). We matched the locations of the same beads by the visual pairing and the consistency of the displacement with their neighbors. In most cases, we apply a global displacement to one of the sets to help the identification of the pairs of beads (Fig S1C, bottom). In general, we picked a subset of marker point pairs with a more homogeneous spacing to ensure a larger triangulation size. We perform the hand-picked process twice, which results in two sets of bead spacings from the same cell.

**Marker point projection onto wall outline:** To ensure that the calculations using the fluorescent bead points and wall outline were done on the same surface, we extrapolated the beads to the corresponding closest point from the cell wall outline (Fig S1E). Any extrapolations that went to an inaccurate (more than 5 pixels) Z-plane of the outline were removed. Additional visual inspection of the projection was done to ensure no projection inaccuracies. The same process was done for the turgid and unturgid beads and outline.

**Automated triangulation additional details:** After applying the automated triangulation scheme detailed in the Methods section of the main text, we would use visual inspection of the triangulations to determine which ones were not accurate to the cell surface. For example, due to limited bead positions at the top of the cell, the automated process would create triangles on the top cutting across the cell.

### 1.3 Computation of curvatures from the wall surface:

Our curvature calculation relies on finite differences to calculate the local tangent vectors. To parameterize the surface regions into a grid, $X(u,v) = (x,y,z)$, we use the Matlab functions `meshgrid` and `griddata` [6, 7]. The grid parametrization can be $(x,y)$, $(x,z)$, or $(y,z)$ depending on the cell surface orientation (Fig 2D). The first function, `meshgrid`, takes in the coordinate range of the parameterized surface along with a specified pixel spacing, which we will discuss at the end of the section. This then returns the corresponding uniform 2D grid coordinates in the parametric space (Fig 2E). The second function, `griddata` with option "`cubic`", will interpolate the original outline at the query points defined by the 2D grid to give us $X(u,v)$ where $u, v$ live on the grid (Fig 2E, F). $X(u,v)$ is then used to calculate the local Weingarten Map [8]:

$$W = \begin{bmatrix} E & F \\ F & G \end{bmatrix}^{-1} \begin{bmatrix} L & M \\ M & N \end{bmatrix} \tag{1}$$

with the first and second fundamental coefficients,

$$\begin{aligned} E &= X_u \cdot X_u \\ F &= X_u \cdot X_v \\ G &= X_v \cdot X_v \end{aligned} \tag{2}$$

$$\begin{aligned} L &= X_{uu} \cdot \hat{n} \\ M &= X_{uv} \cdot \hat{n} \\ N &= X_{uu} \cdot \hat{n} \end{aligned} \tag{3}$$

where $\hat{n}$ is the unit normal on each point and $X_\square = [x_\square, y_\square, z_\square]$. The subscripts, $\square$, are the partial derivatives of $X$ in the $u$ and $v$ directions (Fig 2E, $u, v$ arrows), and those partial derivatives are approximated by the `gradient` function in Matlab [9]. The eigenvalues of $W$ are the principal curvatures, $\bar{\kappa}_1(u,v)$ and $\bar{\kappa}_2(u,v)$ in the parameterized space. Say $\begin{bmatrix} u'_1 \\ v'_1 \end{bmatrix}, \begin{bmatrix} u'_2 \\ v'_2 \end{bmatrix}$ are the eigenvectors of $W$ corresponding to the eigenvalues. Then the principal directions in 3D are: $\vec{\alpha}_1 = u'_1 X_u + v'_1 X_v$ and $\vec{\alpha}_1 = u'_2 X_u + v'_2 X_v$ followed by normalization.

The high resolution of the image raw data introduces local waviness and distortion of the cell wall not relevant to the curvature at the cell size level (Fig S4A). Additionally, we observed that the finite difference method is sensitive at the edges of the grid. We adopted a border removal process to effectively remove outliers along the edges. To filter out the local waviness, we found that increasing the pixel spacing used to create $X(u,v)$ could smooth the results (Fig S4A, B). As the spacing increases, the border removal process will naturally remove larger chunks of data. Although these areas can be recovered with additional parameterizations, for simplicity we have opted to use a spacing of 3 pixels (Fig 2E). Our final principal curvature values include repeating the entire process 8 times by shifting the interpolated mesh. This allows us to cover the whole 1-pixel grid space (Fig 2E, F).

### 1.4 Triangle error analysis

In the main text, $\mathbf{F}$ describes the area deformation of a triangle patch in 3D. To simplify the calculations, we study $\mathbf{F}_{2D} = \mathbf{B}_{2D}\mathbf{A}_{2D}^{-1}$ and how the noise, $\delta$, on $\mathbf{B}_{2D}$ and $\mathbf{A}_{2D}$ affect $\mathbf{F}_{2D}$, measured by $\frac{\|\delta\mathbf{F}\|_2}{\|\mathbf{F}\|_2}$. We obtained the 2D sub-matrices by rotating $\mathbf{A}$ and $\mathbf{B}$ using a rotation matrix [10]. We first assume that the noise acts on each matrix component with $|\delta_{ij}^{a,b}| < \delta$. These are added to the matrices $\mathbf{A}_{2D}$ and $\mathbf{B}_{2D}$, with components $x_{ij}^{a,b}$:

$$\mathbf{A}_{2D} = \begin{bmatrix} \delta_{11}^a & \delta_{12}^a \\ \delta_{21}^a & \delta_{22}^a \end{bmatrix} + \begin{bmatrix} x_{11}^a & x_{12}^a \\ x_{21}^a & x_{22}^a \end{bmatrix}$$

$$\mathbf{B}_{2D} = \begin{bmatrix} \delta_{11}^b & \delta_{12}^b \\ \delta_{21}^b & \delta_{22}^b \end{bmatrix} + \begin{bmatrix} x_{11}^b & x_{12}^b \\ x_{21}^b & x_{22}^b \end{bmatrix}$$

The resulting deformation matrix can then be written as:

$$\mathbf{F}_{2D} = \mathbf{B}_{2D}\mathbf{A}_{2D}^{-1} = \left( \begin{bmatrix} \delta_{11}^b & \delta_{12}^b \\ \delta_{21}^b & \delta_{22}^b \end{bmatrix} + \begin{bmatrix} x_{11}^b & x_{12}^b \\ x_{21}^b & x_{22}^b \end{bmatrix} \right) \cdot \frac{1}{\det(\mathbf{A}_{2D})} \left( \begin{bmatrix} \delta_{22}^a & -\delta_{12}^a \\ -\delta_{21}^a & \delta_{11}^a \end{bmatrix} + \begin{bmatrix} x_{22}^a & -x_{12}^a \\ -x_{21}^a & x_{11}^a \end{bmatrix} \right). \tag{4}$$

We perform a linear perturbation analysis on Eq 4 to obtain the first order approximation of the maximum error bound for $\|\mathbf{F}_{2D} - \mathbf{F}_{2D}^0\|$:

$$\|\mathbf{F}_{2D} - \mathbf{F}_{2D}^0\|_2 \leq \left( \frac{2\|\mathbf{B}_{2D}\|_2}{\det(\mathbf{A}_{2D})} + \frac{\|\mathbf{F}_{2D}^0\|_2}{\det(\mathbf{A}_{2D})}(|x_{11}^a| + |x_{22}^a| + |x_{12}^a| + |x_{21}^a|) + 2\|\mathbf{A}_{2D}^{-1}\|_2 \right) |\delta|. \tag{5}$$

Lastly, we divide by the norm of the deformation to get the relative error:

$$\frac{\|\delta\mathbf{F}\|_2}{\|\mathbf{F}_{2D}^0\|_2} \leq \left[ \frac{2\|\mathbf{B}_{2D}\|_2}{\det(\mathbf{A}_{2D}) \cdot \|\mathbf{F}_{2D}^0\|_2} + \frac{(|x_{11}^a| + |x_{22}^a| + |x_{12}^a| + |x_{21}^a|)}{\det(\mathbf{A}_{2D})} + \frac{2\|\mathbf{A}_{2D}^{-1}\|_2}{\|\mathbf{F}_{2D}^0\|_2} \right] |\delta| = \Gamma|\delta|. \tag{6}$$

The final $\Gamma$ is a function of parameters related to the layout of the triangle. This function describes the sensitivity of the triangle vertices to perturbations from $|\delta|$. Since $|\delta|$ sets the scale of perturbation, we used a preliminary value of $|\delta| = 1$. We found that the distribution of each triangles $\Gamma$ values followed a log-normal distribution. Therefore, we discarded any triangles with a log relative error, $\log(\Gamma)$, one standard deviation above the mean.

### 1.5 Experiment noise estimation

Our measurement relies on the tracking of marker points between the turgid and unturgid configuration. To replicate the noise coming from movement during the switch between these two configurations, we tracked the marker point movement on the turgid cell with no treatment. To obtain an estimate of this we first imaged the fluorescent beads in the turgid configuration as normal. Then, instead of switching the media to the high molarity media, we switched it to the same media to replicate any disturbance in the chamber from the change in flow. After readjusting the center plane of each cell as we would normally, we imaged the cell again. With the two successive images of the same cell, we applied our software to both sets of images using the same parameters to extract the fluorescent bead locations. We selected two sets of beads with one-to-one correspondence between the first image and second image, which we denote $V_b$ and $V_c$, respectively. We then fit a transformation of the second set of beads onto the first using least squares to minimize the function: $\sum(Q \cdot V_c + c - V_b)^2$, where $Q$ is a rotation matrix and $c$ is the translation. Using the optimized $Q$ and $c$, we translated $V_c$ back to $V_b$ and then measured the displacement between the two. The results of this displacement normalized by the minimum cell width are shown in Fig 3D.

### 1.6 Range and distribution of elastic moduli inputs to generate synthetic cells:

The computational membrane deformation model established in [11] can be employed to generate the turgid configuration given the unturgid configuration and specified distribution of elastic moduli. The range of $\bar{K}_h$ and $\bar{\mu}_h$ starting values were based on preliminary results of $\lambda_\theta$ at the cell side, ranging between $1.05 \leq \lambda_\theta \leq 1.25$ [11]. We created three cases of bulk modulus distributions as a function of the cell length, $z$. These include nonlinearly decreasing and linearly decreasing towards the cell tip $z_{tip}$, and a constant distribution. The constant case is simply:

$$\bar{K}_h(z) \equiv \bar{K}_{side} = \bar{K}_{tip} \tag{7}$$

The linear case is:

$$\bar{K}_h(z) = \begin{cases} \bar{K}_{side} & 0 \le z < z_1 \\ \bar{K}_{tip} + (\frac{1}{z_{tip}-z_1})(\bar{K}_{tip} - \bar{K}_{side})(z - z_{tip}) & z \ge z_1 \end{cases} \quad (8)$$

and the nonlinear distribution is given by:

$$\bar{K}_h(z) = \frac{1}{2}(\bar{K}_{side} - \bar{K}_{tip})(1 - \tanh(\frac{z - z_2}{w})) + \bar{K}_{tip} \quad (9)$$

In both cases, $z_1$ and $z_2$ specify where the linear and nonlinear cases begin to decrease towards the tip, respectively. In our case, we used $z_{tip} = 4$, $z_1 = 2$, $z_2 = 3$, and $w = 0.2$ (see Fig 3Bi for cell orientation and $z_{tip}$). In the results, the distributions are converted to the local angle $\alpha$. The steepness of the gradient $\bar{K}_{side}/\bar{K}_{tip}$ was also chosen to reflect the proper elastic stretch ratios and wall outlines. We found that, unlike previously reported [12, 13], a gradient too large would cause non-monotone outlines (see also [14]). Although these morphologies may exist, for the purposes of this analysis and our estimates of *P. patens* tip cell morphologies, the gradient of $\bar{K}_h$ will be limited to $\bar{K}_{side}/\bar{K}_{tip} \le 4.67$ throughout all study cases.

We found that the trend of $\bar{\mu}_h$ in comparison to $\bar{K}_h$ had no significant effects on the cell shape (Fig S6). Despite effects on the elastic stretch ratios, since we do not infer $\bar{\mu}_h$ in this paper, we choose one of the three cases. Here, we have chosen $\bar{\mu}_{side} = \bar{K}_{side} - 2$ and $\frac{\bar{K}_{side}}{\bar{K}_{tip}} = \frac{\bar{\mu}_{side}}{\bar{\mu}_{tip}}$, meaning $\bar{\mu}_{tip} = \frac{\bar{\mu}_{side}\bar{K}_{tip}}{\bar{K}_{side}}$. This ensures a consistent ratio of $\bar{K}_h$ and $\bar{\mu}_h$ across the cell, leading to a constant Poisson's ratio. With $\bar{\mu}_{side}$ and $\bar{\mu}_{tip}$, the distribution of $\bar{\mu}_h$ is given the same distribution as $\bar{K}_h$, calculated by Eqs 7, 8, or 9.

### 1.7 Data integration and multiple cell averaging

**Local angle calculation:** Instead of mapping the data along the length of the cell, we calculated the local angle using the wall outline point's normal $\hat{n}$ and the long axis $\hat{z}$.

$$\alpha = \cos^{-1}(\hat{n} \cdot \hat{z}) \quad (10)$$

Specifically, we used the wall outline of the unturgid configuration. We found that if the turgid outline was non-monotone, our local angle denotation would not be a function of the position along the length of the cell. To avoid mislabeling with the local angle, we used the unturgid outline which was guaranteed to have a monotonically decreasing width. We then used $\alpha$ as a spatial variable to identify a cell wall surface point at different locations and across cells (Fig S3A). This $\alpha$ term can be applied to each cell wall outline point and to each triangle. To locate each triangle, we assigned each triangle to a range of $\alpha$ values (Fig S3B, C). Using the MATLAB `inpolygon` function, we isolated the outline points associated to each triangle to find the $\alpha$ range for each triangle.

**Binning, weight calculation, and multiple cell averaging:** We have denoted our binning range to be $0 \le \alpha \le 90$ degrees with adaptive bin sizes. Due to the lower number of triangles at the tip, our last bin has a range of $0 \le \alpha \le 20$ degrees. Likewise, because of the large amount of data at the side and possible sharp transition of gradient, we split our bins into a range of 5 degrees starting from $70 \le \alpha \le 90$. Every bin between the side and tip has a size of 10 degrees. For the data on each triangle, we sort each triangle into a bin if the triangle covers a part of its range (Fig S2C, right). Therefore, a triangle's data could exist within multiple bins. Within each bin, a weighted mean and standard deviation of the parameter being measured is taken. The weights are determined by the area each triangle occupies in the angle range of the bin. The total area within each bin is used as a weight when calculating the relative error of the inference against the ground truth. Outliers are removed before binning using the `isoutlier` function in MATLAB with the `movemedian` setting with a window size of 10. The data is first sorted by angle, then outlier removal is applied before binning. When averaging multiple cells, we took their mean result in each bin. Then, we took a weighted mean of each cell's mean using the total area of triangles in that bin as the weight for each cell. The standard deviation was also weighted.

**Trial analysis:** For Figs S9-S13, we generated 100 cells for each ground truth. Each cell was given a random set of marker points and therefore unique triangulation. To map the cell number versus error, we randomly picked $n = 1 : 50$ cells from the set of 100. For each set of $n$ cells, their results were averaged and then the error was calculated. This was then repeated 100 times to find the mean and standard deviation of the error at each cell number used. For Fig 7, we randomly picked the standard $n = 10$ cells to average. To obtain a smooth observable trend of the endpoint ratio level sets, we repeated the process 50 times to eliminate the random effect. The trends of the averaged error and endpoint ratio error in Fig 8 are obtained from the same repeated processes.

**Multiple Triangulations:** For cell inferences with two triangulations, we created another set of marker points by randomly drawing from the overall set. From there, we could create another triangulation on the same cell and measure the parameters. We then combined the two triangulation data at the binning step to obtain one inference result for the single cell.
